# *Corynebacterium matruchotii* fitness enhancement of adjacent streptococci by multiple mechanisms

**DOI:** 10.1101/2022.05.09.491233

**Authors:** Eric Almeida, Surendra Puri, Subhashini Elangovan, Jiyeon Kim, Matthew Ramsey

**Affiliations:** The University of Rhode Island, Department of Cell and Molecular Biology, Department of Chemistry, Kingston RI 02881; The University of Rhode Island, Department of Chemistry, Kingston RI 02881

## Abstract

Polymicrobial biofilms are present in many environments particularly in the human oral cavity where they can prevent or facilitate the onset of disease. While recent advances have provided a clear picture of both the constituents and their biogeographical arrangement, it is still unclear what mechanisms of interaction occur between individual species in close proximity within these communities. In this study we investigated two mechanisms of interaction between the highly abundant supragingival plaque (SUPP) commensal *Corynebacterium matruchotii* and *Streptococcus mitis* which are directly adjacent *in vivo*. We discovered that *C. matruchotii* enhanced the fitness of streptococci dependent on its ability to detoxify streptococcal-produced hydrogen peroxide and its ability to oxidize lactate also produced by streptococci. We demonstrate that the fitness of adjacent streptococci was linked to that of *C. matruchotii* and that these mechanisms support the previously described “corncob” arrangement between these species but that this is favorable only in aerobic conditions. Further we utilized scanning electrochemical microscopy (SECM) to quantify lactate production and consumption between individual bacterial cells for the 1^st^ time, revealing that lactate oxidation provides a fitness benefit to *S. mitis* and not pH mitigation. This study describes mechanistic interactions between two highly abundant human commensals that can explain their observed *in vivo* spatial arrangements and suggest a way by which they may help preserve a healthy oral bacterial community.

## Introduction

Over the past decades our knowledge of the human oral microbiome has increased drastically, revealing a robust polymicrobial biofilm in supragingival plaque (SUPP) that is present in healthy as well as diseased conditions. While we know a great deal about what bacteria reside in SUPP (Benítez-Páez et al., 2014; Eren et al., 2014; Schoilew et al., 2019; Xiao et al., 2016), we know very little about the interactions between taxa especially in healthy conditions relative to disease. Given that dysbiosis of the healthy microbiota is often a prelude to oral disease, we wish to study interactions within the healthy community to potentially reveal any community members that might help preserve stable community structure and constituency, potentially preventing the onset of disease.

Previous studies have shown the importance of attachment to the development of the oral biofilm (Kolenbrander et al., 2006) and new data has identified and refined the spatial organization of abundant commensal organisms found in SUPP (Mark Welch et al., 2016). Human microbiome project (HMP) data and recent microscopy of healthy individuals has revealed that one of the most abundant and prevalent species in SUPP is *Corynebacterium matruchotii* (Mark Welch et al., 2016; Schoilew et al., 2019; Xiao et al., 2016). It has been correlated with good dental health and hypothesized to be important in the organization of some plaque biofilm structures particularly due to its ability to adhere to *Streptococcus* species forming a structure referred to as a “corncob” with the *Corynebacterium* filament being surrounded by streptococci (Mark Welch et al., 2016), a role typically ascribed to *Fusobacterium* (Foster and Kolenbrander, 2004; Kolenbrander et al., 2006). Also, *C. matruchotii* has shown *in vitro* to be able to co-aggregate with *Actinomyces* species which are known to be early colonizers during the plaque biofilm formation (Esberg et al., 2020). It’s been hypothesized that *C. matruchotii* binds to an existing biofilm of *Streptococcus* and *Actinomyces* cells for attachment and anchoring to the plaque (Esberg et al., 2020; Mark Welch et al., 2016). The spatial organization of microbes in SUPP has been characterized in the ‘hedgehog’ model (Mark Welch et al., 2016) which visualizes *C. matruchotii* and its proximity to adjacent *Streptococcus* species, such as *S. mitis* at the SUPP perimeter (Mark Welch et al., 2016; Morillo-lopez et al., 2021).

*Streptococcus* species, such as *S. mitis*, are one of the most abundant species in the oral microbiome (Eren et al., 2014). *Streptococcus* species have deployed many tactics to compete in their environment such as producing antimicrobial metabolites like H_2_O_2_ (Redanz et al., 2018) through the activity of pyruvate oxidase (*spxB*) which takes pyruvate, phosphate, and molecular oxygen and converts them into acetyl phosphate, CO_2_, and H_2_O_2_ (Abranches et al., 2018). This production of H_2_O_2_ has been shown to affect the oral community composition and it is hypothesized in this community *Streptococcus* species metabolize oxygen and sugars to produce H_2_O_2_ and lactate while attached to *C. matruchotii* (Zhu and Kreth, 2012). Other streptococci are reported to co-aggregate with catalase positive organisms to benefit from catalase activity (Jakubovics et al., 2008) and crossfeeding on *Streptococcus-*produced lactate by commensal microbes which enhances their yield has been previously shown (Ramsey et al., 2011). Others have previously hypothesized that *C. matruchotii* and adjacent streptococci would also utilize these same mechanisms in *in vivo* hedgehog structures (Mark Welch et al., 2016). *C. matruchotii* is in close proximity to many streptococci but not much is known about how it endures these stressors. At low pH streptococci can create an environment suitable for the pathogenic *Streptococcus mutans* to thrive in the community and cause caries (Kim et al., 2020; Takahashi, 2005; Van Houte et al., 1991). If interactions between *C. matruchotii* and *S. mitis* create a more stable and healthy plaque formation without enhancing accumulation of H_2_O_2_ or creation of low pH environments, this relationship could strengthen the plaque’s pathogen excluding properties known as colonization resistance (Abt and Pamer, 2014). Despite their co-proximity and abundance in these structures, little is known about how these organisms interact (Mark Welch et al., 2016). Previous studies have not likely appreciated the role *C. matruchotii* plays in bridging early and late colonizers within the plaque (Foster and Kolenbrander, 2004; Kolenbrander et al., 2006) and its importance in the structuring of the plaque community. *C. matruchotii*, in close proximity with *S. mitis*, faces the task of detoxifying the streptococcal-produced metabolites being excreted into the environment. This paper focuses on how *C. matruchotii* interact biochemically with H_2_O_2_ and lactate produced by *S. mitis*.

We employed a reductionist approach to investigate the relationship between these species in *in vitro* biofilms. We discovered that *S. mitis* obtained a significant increase in growth yield with *C. matruchotii* aerobically and this growth benefit is lost anaerobically where *C. matruchotii* growth is inhibited by *S. mitis*. Likewise, when *C. matruchotii* oxidative stress responses were altered its fitness and the coculture benefit to *S. mitis* yield were reduced. We also observed that *C. matruchotii* upregulated lactate catabolism genes when in close proximity with *S. mitis*. Removal of streptococcal-produced lactate by *C. matruchotii* was a contributor to *S. mitis* growth benefit and surprisingly this effect was pH-independent. We also utilized scanning electrochemical microscopy to demonstrate that lactate catabolism can deplete local concentrations of this organic acid swiftly in real-time at sub-micron scales, implying that acid removal in coculture can occur in observed *in vivo* arrangements between these organisms. These data suggests *C. matruchotii* has the ability to maintain *S. mitis* growth in SUPP by aiding in detoxifying reactive oxygen species (ROS) and removing streptococci-produced lactate which likely helps preserve a robust polymicrobial biofilm *in vivo*.

## Materials and Methods

### Strains and media

Strains and plasmids used in this study are listed in Table S1. *C. matruchotii* (ATCC 14266) and *S. mitis* (ATCC 49456) were grown on Brain Heart Infusion media supplemented with 0.5% yeast extract (BHI-YE) at 37°C in a static incubator with 5% CO_2_ or in 5% H_2_, 10% CO_2_ and 85% N_2_ in anaerobic conditions. *E. coli* was grown at 37 °C in standard atmospheric conditions with liquid cultures shaken at 200 RPM. Antibiotics were used at the following concentrations: kanamycin 40 µg/ml for *E. coli* and 10 µg/ml for *C. matruchotii*.

### Colony biofilm coculture/ buffered coculture/ catalase coculture

Overnight cultures of *C. matruchotii* and *S. mitis* species were grown in Brain Heart Infusion media supplemented with 0.5% yeast extract (BHI-YE) at 37°C in a static incubator with 5% CO_2_ or in 5% H_2_, 10% CO_2_ and 85% N_2_ for anaerobic conditions. Colony biofilm assays were carried out as described previously (Merritt et al., 2005). Briefly, A semi permeable 0.22µm polycarbonate membrane filter (Zheng and Stewart, 2002) was placed on solid BHI-YE media (supplemented with 1.6% agar). Ten µl of each culture were spotted on the membrane filters and monocultures were spot with 10µL of BHIYE. The cultures incubated for 48hr and the membranes were placed in a microcentrifuge tube with 1mL of BHI-YE. The tubes were vortexed to resuspend into media and serially diluted and track plated (Jett et al., 1997) to count colony forming units per mL (CFU/mL). *S. mitis* was counted by using BHI-YE plates and *C. matruchotii* on BHI-YE plates supplemented with 100 µg/ml fosfomycin. Buffered and pH indicator cocultures were carried out with 50mM MOPS and 18mg/mL of phenol red added to BHI-YE. Catalase cocultures were carried out with 100U/mL of catalase added to BHI-YE.

### RNAseq experiment and analysis

Mono and cocultures were prepared similar to the colony biofilm coculture with the exception that culture membranes were incubated for 24hr and moved to fresh media for an additional 4hr. Membranes were then removed from solid agar and immediately placed into RNALater (Ambion) where cells were removed by agitation and pelleted by centrifugation. Cell pellets were stored in Trizol reagent at -80°C. Experiments were carried out in biological duplicates. RNA extraction, library preparation and sequencing were then carried out by the Microbial ‘Omics Core facility at the Broad Institute. RNASeq libraries were generated using previously described methods (Shishkin et al., 2015). Sequence data was aligned using Bowtie2 (Langmead and Salzberg, 2012) and read counts per coding sequence were called using HTSeq-Count (Anders et al., 2015). Statistical analysis was carried out via DESeq2 (Love et al., 2014) to determine differentially expressed genes. Scripts of this pipeline can be found at https://github.com/dasithperera-hub/RNASeq-analysis-toolkit. Sequence libraries are available through the NCBI short read archive (SRA) under bioproject number PRJNA832032.

### Gene deletions

All *C. matruchotii* gene deletions were carried out with sucrose counterselection using a suicide vector derived from pMRKO (Ramsey et al., 2011), pEAKO2 which contains *sacB* from pK19mobsacB (Schgfer et al., 1994). Approximately 1000 bp up and downstream flanking regions for each gene were used for homologous recombination and fragments were cloned into pEAKO2 via Gibson Assembly (Gibson et al., 2009). *C. matruchotii* cells will be made as previously described (Takayama et al., 2003). Transformations were carried out with 50µL of competent cells and 1µg of DNA electroporated with 0.2cm gap cuvettes at 2.5kV voltage, 400Ω resistance, and 25µF capacitance. After electroporation, 950mL of prewarmed BHI-YE will be added to the cuvette and the mixture will be moved to a 46°C heat block for 6 minutes. After heat shock, transformations will be shaken at 250 RPM at 37°C for 4hr. Transformations were plated on BHIYE Kan_10_ plates and incubated for 4 days at 37°C. Mutants were verified through PCR.

### Limiting glucose coculture

Cultures were prepared like colony biofilm coculture described above except for being inoculated into 2mL of liquid defined medium. Modified RPMI medium (Gibco) was used as a base and supplemented with 8mM glucose. Cocultures were inoculated for 48 hrs and track plated to determine CFU/ mL.

### Growth curves

Five milliliter cultures of *C. matruchotii* were grown in BHI-YE for 48hr and back diluted in modified RPMI media supplemented with 40mM lactate at an OD_600_ of 0.025. Cultures were incubated statically at 37°C with 5% CO_2_. Optical density readings were taken every 4hr over a 72hr period.

### Scanning electrochemical microscopy sample preparation

Bacteria were grown overnight in BHI-YE and then washed by centrifugation in defined medium. The defined medium used was an amended version of Teknova EZ RICH (Teknova, M2105). We prepared the medium as described in the manufacturers instructions with the addition of vitamin solution, lipoic acid, folic acid, riboflavin, NAD+ and nucleotides to final concentrations from that of an oral complete defined medium previously described (Brown and Whiteley, 2007) and glucose at 10 mM. Bacteria were grown to an OD600 of 0.5-0.9 and then diluted to final concentrations of 1.2×107 and 6.0×106 CFU of *C. matruchotii* and *S. mitis* respectively in 200 µL of defined medium. This was then incubated at 37°C with 5% CO_2_ for 1 h. 10 µL of this solution was then added to a poly-lysine coated glass slide and incubated at 37°C for 15 minutes after which medium was removed by micropipette to remove planktonic cells and ensure only attached cells remained. An additional 10uL of prewarmed defined medium was then added to the slide. Samples were then transferred to the SECM instrument for further analysis.

### Scanning electrochemical microscopy acquisition

Scanning parameters and nano-probe design are similar to methods described previously (Connell et al., 2014; Kim et al., 2014; Kim et al., 2016; Puri and Kim, 2019). A full description of SECM calibration, sample acquisition and metabolite quantification are provided in the Supplemental Materials.

## Results

### *C. matruchotii* enhances the growth of *S. mitis* in aerobic conditions

We performed pairwise coculture experiments aerobically and anaerobically with a colony biofilm model (Merritt et al., 2005) on solid medium to quantify growth yield differences between mono- and cocultures of *S. mitis* with *C. matruchotii* (Fig. 1). Using this reductionist approach, we observed a 954-fold increase in growth yield of *S. mitis* in coculture. Unexpectedly, *C. matruchotii* had no significant difference in growth yield with *S. mitis* (Fig. 1A). While previous studies have hypothesized that *C. matruchotii* – *Streptococcus* interactions occur in aerobic microenvironments within SUPP (Mark Welch et al., 2016; Morillo-lopez et al., 2021), we also performed the same experiment in anaerobic conditions as a comparison (Fig. 1B). Interestingly, the coculture growth benefit for *S. mitis* was lost while *C. matruchotii* growth yield decreased ∼130-fold. To investigate how *C. matruchotii* enhances *S. mitis* growth yield in coculture we performed RNAseq to compare mono-vs coculture transcriptome data.

**FIG 1.**
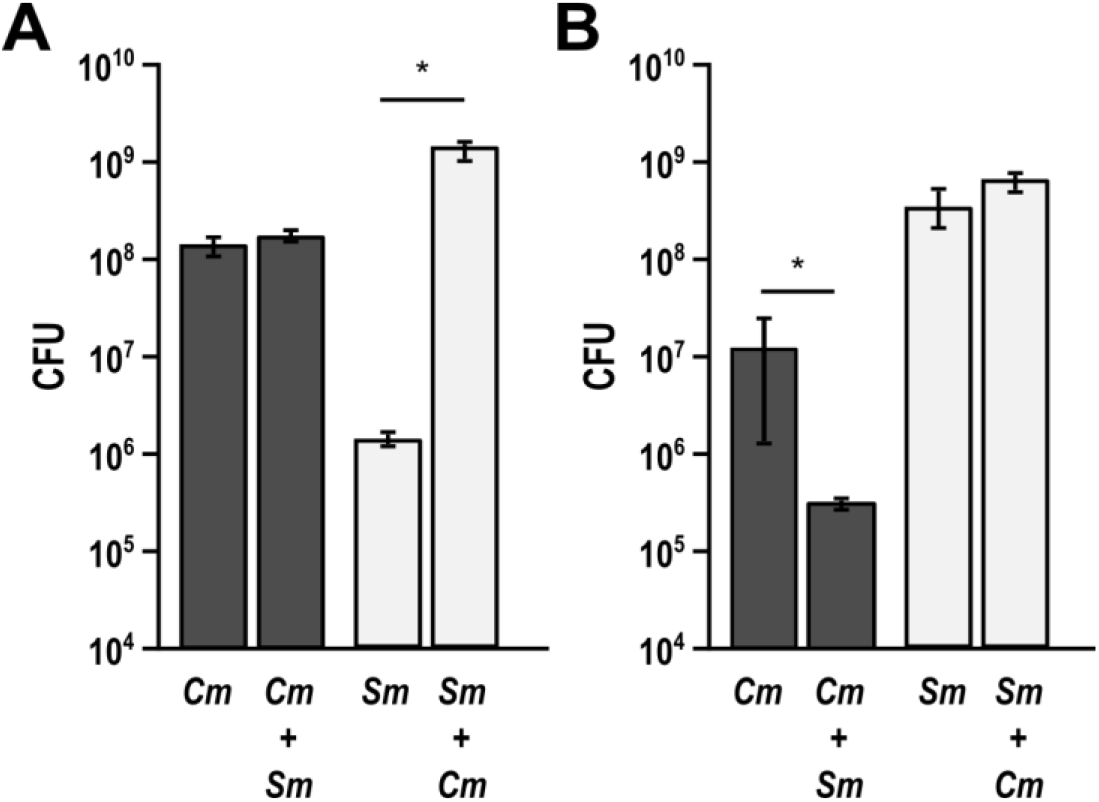
Growth yield measurements of mono vs coculture biofilms. Aerobic (A) and anaerobic (B) CFU counts of *C. matruchotii* (*Cm*) and *S. mitis* (*Sm*) in mono and cocultures. Data are mean CFU counts for n≥3 and error bars represent 1 standard deviation. * denotes p< 0.05 using a Student’s t-test.

### *C. matruchotii* upregulates genes necessary for L-lactate catabolism and oxidative stress response

*C. matruchotii* differentially expressed only 22 genes (greater than 2-fold) in aerobic coculture with *S. mitis* (Table S2). Interestingly, C. *matruchotii* upregulated the *lutABC* operon (*lutA*, 4.37fold; *lutB*, 3.76-fold; *lutC*, 3.20-fold), whose gene products in *Bacillus subtilis* catabolize L-lactate (Chai et al., 2009) using oxygen as a terminal electron acceptor; therefore, in the absence of oxygen, *C. matruchotii* is no longer able to catabolize L-lactate, as previously shown (Iwami et al., 1972). *C. matruchotii* also significantly upregulates a bacterial non-heme ferritin-encoding gene (2.39-fold) in coculture. This protein has been characterized in *Mycobacterium smegmatis* to sequester ferrous ions as part of the oxidative stress response (Smith, 2004). Given the coculture growth and transcriptome results, we broadly hypothesized that *C. matruchotii* crossfeeds on *S. mitis*-produced lactate while detoxifying *S. mitis*-produced H_2_O_2_ similar to other microbes in the oral cavity (Jakubovics et al., 2008; Ramsey et al., 2011). Given the fact that *C. matruchotii* cannot utilize L-lactate anaerobically and *S. mitis* is only provided a growth benefit in the presence of oxygen, we believe these data suggest one mechanism by which the biogeography of these species *in vivo* could be influenced by their metabolic interactions.

### Lactate utilization by *C. matruchotii* influences *S. mitis* growth yield

The growth enhancement of *S. mitis* in coculture with *C. matruchotii* is likely due to several factors including H_2_O_2_ decomposition and lactate catabolism. It is unclear if the removal of lactate itself or the removal of lactate and subsequent increase in pH is responsible for *S. mitis* growth yield enhancement. We 1^st^ tested the impact of pH by performing growth experiments in the same medium with increased buffer capacity by adding 50 mM MOPS. Qualitatively, we observed that *S. mitis* monoculture colonies no longer produced yellow coloration in buffered medium containing the pH indicator dye phenol red (i.e. no longer acidified the environment) compared to the original medium (data not shown). Quantitatively, we observed that *S. mitis* growth yield had no significant change in monoculture with additional MOPS (Fig. S1) indicating that pH was likely not responsible for *S. mitis* growth yield increases in coculture.

To determine if removal of lactate by *C. matruchotii* via catabolism was enhancing streptococcal fitness we constructed a *lutA* gene deletion mutant (*ΔlutA*) since each gene within the *lutABC* operon had been described to be essential for L-lactate catabolism (Chai et al., 2009). The *ΔlutA* strain was significantly impaired in L-lactate utilization showing a diminished growth rate (doubling times of 17.9h for the wt and 27.4h for *ΔlutA*) and yield aerobically with L-lactate as the sole carbon source (Fig. S2). A full *lutABC* operon deletion strain was also created and showed similar results (data not shown). In *B. subtilis*, each gene within the *lutABC* operon is essential for lactate oxidation and it can no longer use lactate as its sole carbon source (Chai et al., 2009). *C. matruchotii* possesses two additional annotated L-lactate dehydrogenases which may function bidirectionally allowing it to more slowly oxidize L-lactate without a functional *lutABC* system.

We next tested the *ΔlutABC* mutant in mono vs coculture with *S. mitis* to determine if impaired lactate utilization led to a decrease in *S. mitis* yield in coculture with *C. matruchotii*. Using defined medium in glucose-limited conditions to force the bacteria to compete for the limited carbon source and/or promote cross-feeding on streptococcal produced lactate, we performed mono vs cocultures and determined that both *S. mitis* and *C. matruchotii* Δ*lutABC* fitness were significantly decreased in coculture (Fig. 2). *C. matruchotii ΔlutABC* can only poorly catabolize L-lactate and thus poorly cross-feed on *S. mitis*-produced L-lactate compared to the wildtype. As *ΔlutABC* and *S. mitis* are now forced to compete for limited glucose, both exhibit a decreased growth yield. This is in agreement with previous data anaerobically (Fig. 1B), where lactate oxidation by *C. matruchotii* does not occur. The growth yield increase of *S. mitis* in coculture is diminished when *C. matruchotii* cannot oxidize lactate but this does not fully explain the total growth benefit provided, suggesting another mechanism(s) at work.

**FIG 2.**
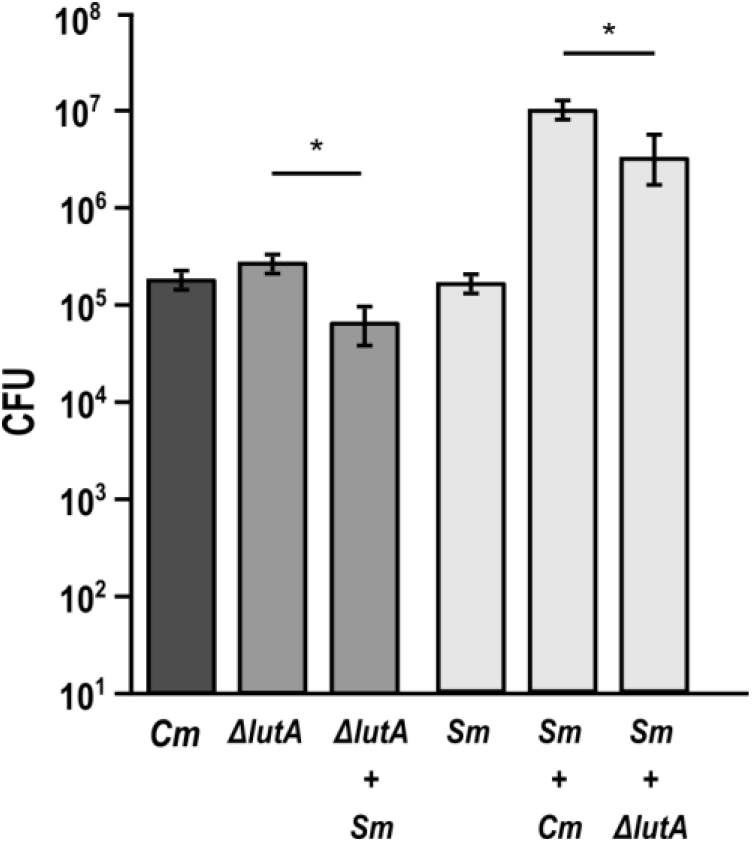
Limiting glucose colony biofilm cocultures. CFU counts of *C. matruchotii* (*Cm*), *C. matruchotii ΔlutA*, and *S. mitis* (*Sm*) in mono and cocultures. Data are mean CFU counts for n≥3 and error bars represent 1 standard deviation. * denotes p< 0.05 using a Student’s t-test.

### Catalase abundance leads to enhanced streptococcal growth yields

Given that lactate oxidation by *C. matruchotii* provides only a small portion of the fitness benefit in coculture to *S. mitis* we next investigated if H_2_O_2_ detoxification by *C. matruchotii* also contributes to fitness. Surprisingly, in coculture with *S. mitis, C. matruchotii* did not upregulate expression of the single catalase (*katA*) encoded on its chromosome. We observed that catalase was already maximally expressed aerobically and not expressed anaerobically (data not shown). To test if catalase-dependent H_2_O_2_ detoxification was important both for *C. matruchotii* fitness in coculture and subsequent *S. mitis* growth yield enhancement, we generated the catalase gene deletion mutant, *C. matruchotii ΔkatA*. Interestingly, this mutant had to be generated entirely under anaerobic conditions and does not survive incubation in aerobic or microaerophilic conditions (data not shown), making it impossible to test this mutant in aerobic coculture with *S. mitis*. Instead, we determined the contribution of catalase to the growth of these species by adding it exogenously. We performed aerobic mono vs cocultures in growth medium amended with 100U/mL of bovine catalase.

Previous studies (Eisenberg, 1973; Jakubovics et al., 2008; Regev-Yochay et al., 2007) have indicated that streptococcal-produced H_2_O_2_ is capable of limiting their own growth. We observed that adding exogenous catalase elevated the monoculture growth yield of *S. mitis* 6.42-fold (Fig. 3A). This self-limitation by H_2_O_2_ production is also observed when comparing monoculture fold changes of *S. mitis* to the non H_2_O_2_–producing *ΔspxB* mutant (Fig. 3). Interestingly, the growth benefit of *S. mitis* in coculture with *C. matruchotii* dropped from 954-fold to 148-fold when amended with catalase.

**FIG 3.**
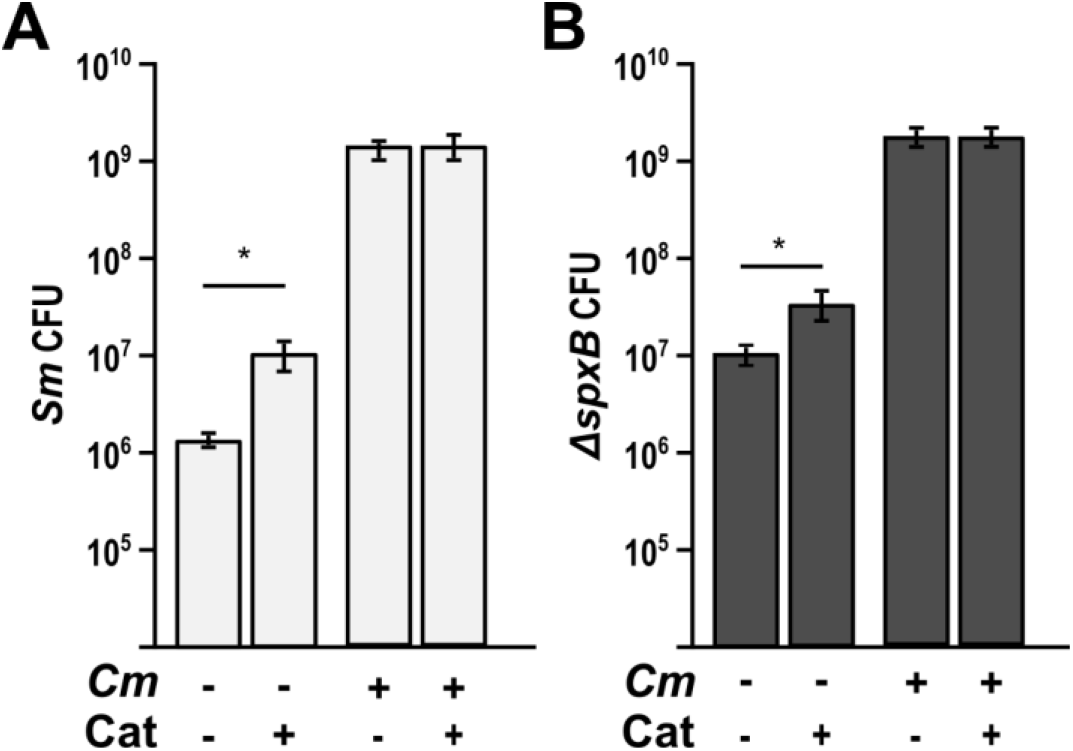
*S. mitis* monoculture enhanced by exogenous catalase. A) CFU counts of *S. mitis* WT (*Sm*) in mono and coculture with *C. matruchotii* (*Cm*) on media containing 100U/mL of catalase vs without. B) CFU counts of *S. mitis ΔspxB* in mono and coculture with *Cm* on media containing 100U/mL of catalase vs without. * denotes p< 0.05 using a Student’s t-test.

### *C. matruchotii* requires a functional oxidative stress response to be fit to interact with *S. mitis*

In coculture with *S. mitis, C matruchotii* significantly upregulated a gene encoding ferritin, a bacterial non-heme protein involved in oxidative stress response (Smith, 2004). We hypothesized that ferritin was needed for *C. matruchotii* fitness with H_2_O_2_-producing streptococci. To test this, we deleted the ferritin encoding gene generating *C. matruchotii Δftn* and performed cocultures with WT *S. mitis* and *S. mitis ΔspxB* (which is unable to produce H_2_O_2_) (Redanz et al., 2018). In coculture with WT *S. mitis*, we observed that the *Δftn* mutant fitness decreased 7.35-fold (Fig. 4A) and this decrease was not observed in coculture with the *S. mitis ΔspxB* strain. *S. mitis* WT had a 4.6-fold significant decrease in growth yield with *C. matruchotii Δftn* compared to *C. matruchotii* WT whereas there was no change in growth yield with *S. mitis ΔspxB* with either *C. matruchotii* strain. This shows that *C. matruchotii* needs a functional oxidative stress response in order to be fit with its interactions with *S. mitis* WT. These data indicate that H_2_O_2_ detoxification is the largest contributor to enhanced *S. mitis* fitness in coculture but also that other mechanisms, likely *C. matruchotii* lactate oxidation, further enhance fitness.

**FIG 4.**
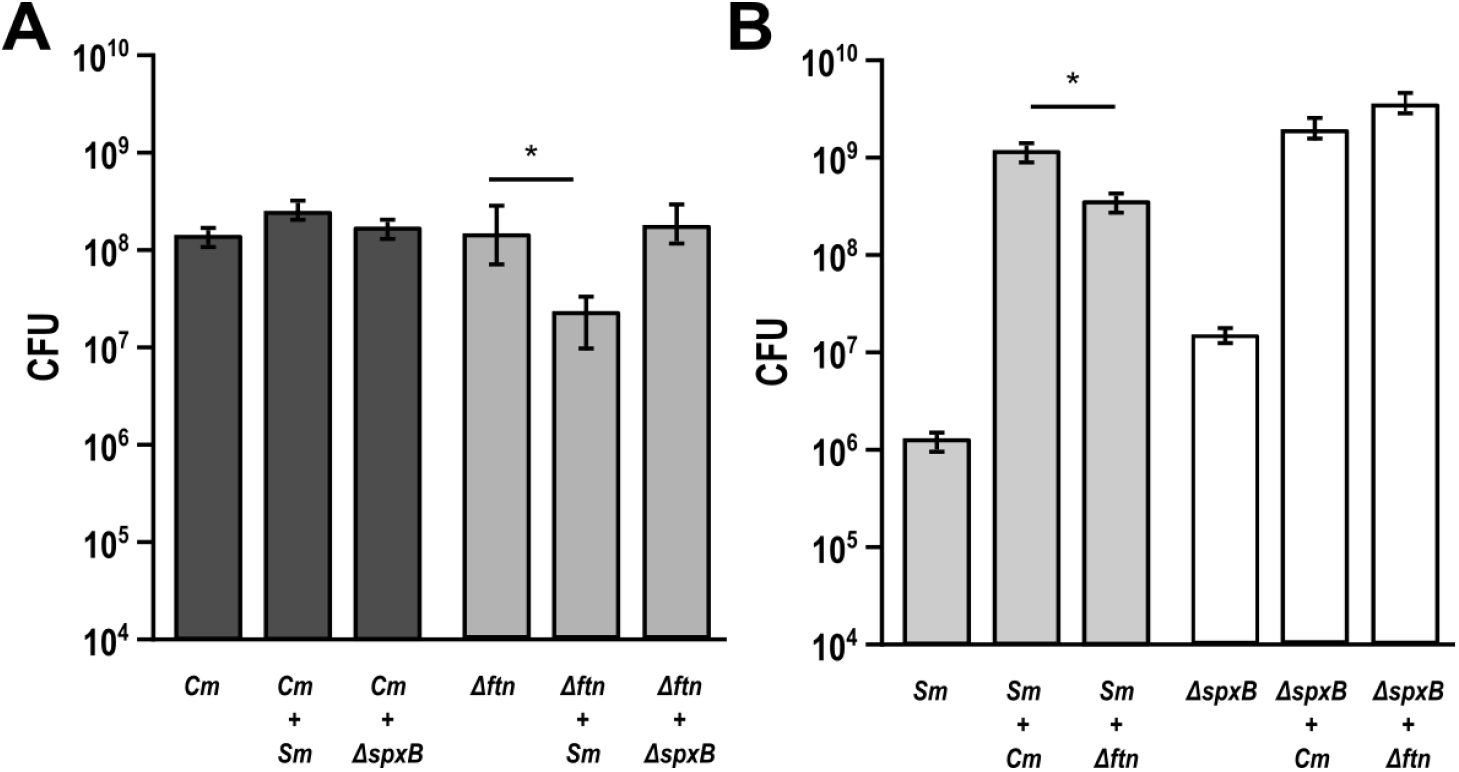
*C. matruchotii* ferritin knockout inhibited when cocultured with *S. mitis*. A) Aerobic CFU counts of *C. matruchotii* WT (*Cm*) and ferritin knockout (*Δftn*) in mono and coculture with *S. mitis* WT and strain lacking ability to create H_2_O_2_ (*ΔspxB*). B) CFU counts of *Sm* and *ΔspxB* in mono and coculture with *Cm* and *Δftn*. Data are mean CFU counts with error bars indicating standard deviation for n≥3. * denotes p< 0.05 using a Student’s t-test compared to monoculture.

### Scanning electrochemical microscopy (SECM) reveals oxidation of *S. mitis*-produced lactate by adjacent *C. matruchotii* at sub-micron scale

To investigate lactate production and consumption *in situ* by bacteria as well as the topography of bacterial cells, a submicropipet-supported interface between two immiscible electrolyte solutions (ITIES) was employed (Figs 5, S3) (Puri and Kim, 2019). With this submicrotip, an etched Ni/Cu electrode in the internal organic electrolyte exerts a bias across the submicroscale liquid/liquid interface against an electrode in the aqueous solution (Fig. S3A) to yield the amperometric tip current based on the selective interfacial transfer of a small probe ion (Puri and Kim, 2019). The coculture of *C. matruchotii* and *S. mitis* was immobilized over a poly L-lysine coated glass plate, and studied by scanning or approaching an 800 nm-diameter pipet tip over the bacteria (Fig. S3C). Further detail is provided in the supplemental materials.

**FIG 5.**
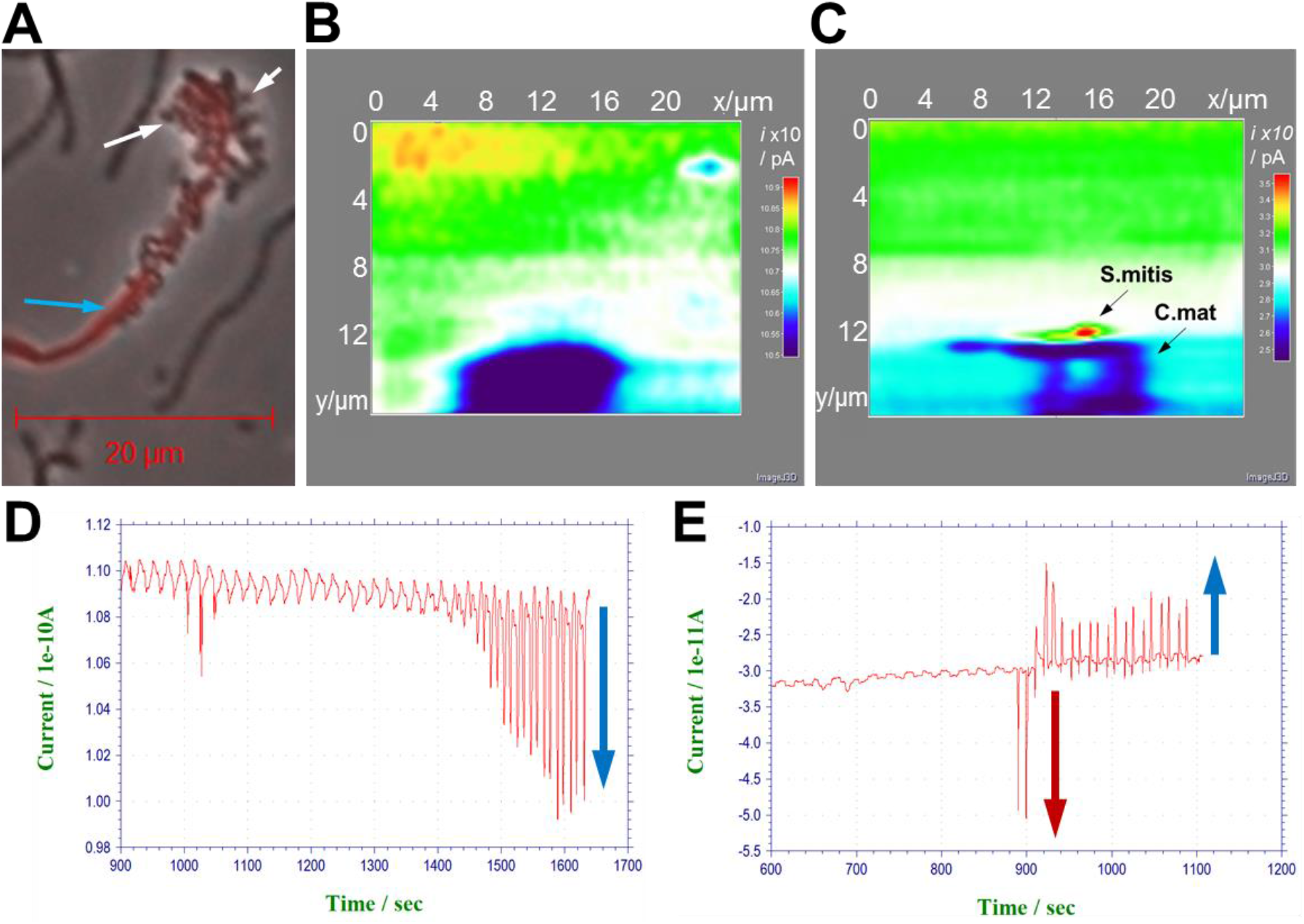
(A) Optical microscopic image of *C. matruchotii* (blue arrow) and *S. mitis* (white arrows) coculture. Constant-height SECM images based on (B) TEA^+^ IT (obtained with a gap between the tip and bacteria, *d*_c_ = 1.8 *d/a*) and (C) lactate IT (obtained at *d*_c_ = 1.2 *d/a*), Chronoamperometric responses based on (D) TEA^+^ IT and (E) lactate IT (raw data, cross sections of SECM images in (B) and (C), respectively). The current polarity is set to positive for cationic current response and negative for anionic current response.

We employed the constant-height mode of submicroscale SECM to successfully image single bacterial cells in coculture (Fig. 5). The high spatial resolution was obtained by using submicropipet tips, which were characterized by cyclic voltammetry for tetraethylammonium (TEA+) ion transfer (IT) in situ to obtain a diffusion limited current in the bulk solution, *i*_T,∞_ (120 pA). The submicrotip approached the glass substrate until the tip current decreased to 90 % of *i*_T,∞_, which is equivalent to the tip-substrate distance, *d*, of 0.85 µm with the tip radius, *a*, of 430 nm. Further, a tip was withdrawn 1.75∼2.00 µm higher, and scanned laterally at the fixed height while the tip current was monitored to obtain an SECM image (Fig. S3D). Constant-height imaging of cocultured bacteria for the probe ion TEA was obtained with the gap between the tip and bacteria, *d*_c_ = 0.75 µm, i.e., 1.80 normalized distance to tip radius, (*d/a*). This SECM image could not resolve each individual bacterial cell. For instance, a lump was identified in 25 µm × 20 µm image based on TEA^+^ IT, which did not resolve any difference between bacterial cells (Fig. 5B). Low tip currents of ∼ 80 % of *i*_T,∞_ for TEA^+^ above these bacteria was obtained due to hindered diffusion of TEA^+^ by adjacent bacteria with membranes near impermeable to this probe ion. As shown in the chronoamperometric responses (shown as raw data in Fig. 5D, cross sections of the SECM image in Fig. 5B), currents were monotonically lowered in 1450∼1650 s (positive polarity for cationic currents).

The same area in 25 µm × 20 µm was imaged based on lactate IT with the gap between the tip and bacteria, *d*_c_ = 0.50 µm (1.20 *d/a*), which could resolve individual *S. mitis* and *C. matruchotii* clearly (Fig. 5C). An initial current, ∼30 pA above a glass substrate corresponds to 0.26 mM of lactate produced by ensemble of bacteria and diffused to bulk solution near bacteria according to eq 1 below.

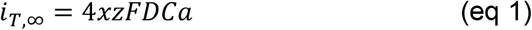

where *i*_T,∞_ is a current in bulk, *x* is the function of RG ratio (RG is the ratio of outer and inner diameters of a glass pipet, 1.16 for a RG 1.5 tip), *z* is charge of lactate, *F* is Faraday constant (96485 C/mol), *D* is the diffusion coefficient (6 × 10^−6^ cm^2^/s), *C* is a concentration of lactate (0.26 mM), and *a* is the inner radius of a pipet tip (430 nm).

In this SECM image, high tip currents, 50 pA of 166 % of *i*_T,∞_ for lactate are obtained above spherical *S. mitis*, while low tip currents, 22 ∼24 pA of 75∼80 % of *i*_T,∞_ are observed above filamentous *C. matruchotii*. Not only distinctive morphologies are clearly distinguished between two different bacteria as shown in optical microscopic image (Fig. 5A), but also the production and consumption of lactate between them are visually confirmed *in situ*. As shown in chronoamperometric responses (shown as raw data in Fig. 5E, cross sections of the SECM image in Fig. 5C), currents were dramatically transposed from an enhanced response over *S. mitis* to reduced ones over *C. matruchotii* (negative polarity for anionic currents), implying local increase in lactate produced by *S. mitis* and local depletion of lactate consumed by *C. matruchotii*. Notably, this SECM image successfully visualized the chemical interaction between two commensal oral microbes in real time and is the 1^st^ SECM study to our knowledge that measures metabolite exchange between two individual bacterial cells. Specifically, *S. mitis* produces ∼0.52 mM lactate locally, which is efficiently depleted by C. *matruchotii* (Figs. 5, S4) thus verifying a standing question about their commensal relationship that cannot be answered only by optical microscopic imaging. Quantitative analysis of the permeability of the bacterial membrane and the local concentration of lactate produced by *S. mitis* are discussed in the supplemental materials.

## Discussion

Interactions between commensal bacteria within healthy SUPP are understudied for their role in maintaining plaque homeostasis and host health compared to subgingival plaque and oral disease (Foster and Kolenbrander, 2004; Guggenheim et al., 2001; Kolenbrander et al., 2006). While the organisms in SUPP are in close proximity to one another and capable of physical and biochemical interaction, these mechanisms are largely hypothetical (Mark Welch et al., 2016). Characterizing the behavior of abundant SUPP commensal organisms can help reveal necessary interactions that could maintain a healthy microbiome. One set of interactions are those between *Corynebacterium matruchotii*, and *Streptococcus* spp in previously described ‘hedgehog’ structures (Mark Welch et al., 2016; Morillo-lopez et al., 2021) where they occur at the presumed aerobic biofilm / saliva margin. This study investigates these interactions and provides novel data on metabolite exchange between individual cells that has broad implications on polymicrobial biofilms beyond the human oral cavity.

Our data indicates that *S. mitis* had a significant growth yield increase when cocultured with *C. matruchotii* (Fig. 1A) and this growth benefit was lost anaerobically (Fig. 1B) which aligns with their proximal association only at the aerobic margin of their biofilm structures (Mark Welch et al., 2016; Morillo-lopez et al., 2021). H_2_O_2_-producing *Streptococcus* and adjacent commensal species have been shown to coexist despite ROS production (Jakubovics et al., 2008). *C. matruchotii* is uninhibited when cocultured with *S. mitis* aerobically likely due to catalase production. A *C. matruchotii ΔkatA* was created to test this hypothesis and unexpectedly would only grow anaerobically. To confront this limitation and understand the role catalase has on *S. mitis* growth benefit, we added exogenous catalase into the medium to observe if there was a less prominent or loss of growth benefit between mono and coculture *S. mitis* since both now benefit from detoxified ROS. *S. mitis* monocultures on media containing catalase showed increased growth yield (Fig. 3A), confirming the role of catalase in enhancing *S. mitis* fitness. However, even with exogenous catalase there was still a significant increase in *S. mitis* yield in coculture suggesting that *C. matruchotii* provides further growth benefits beyond ROS detoxification.

The fitness of *C. matruchotii* in *S. mitis-*induced oxidative stress is influenced by its ability to not only detoxify H_2_O_2_ but prevent its reaction with free ferrous ions. This was evident by the expression of a ferritin-like protein in coculture with *S. mitis* that has an 82% protein identity with the ferritin encoded in *Corynebacterium mustelae*. Bacterial ferritin-like proteins have been shown to sequester away iron to avoid the oxidation of ferrous iron to ferric iron (Smith, 2004). Ferritin-like proteins can also bind to DNA for protection from these free hydroxyl radicals (Smith, 2004). Their activity can prevent the production of hydroxyl radicals known to damage DNA and lipids (Winterbourn, 1995). *C. matruchotii Δftn* showed a significant yield decrease in coculture with *S. mitis* but this inhibition was not observed with *S. mitis ΔspxB*. We observed that any decreases in *C. matruchotii* yield were mirrored by decreases in *S. mitis* yield as well, linking streptococcal fitness to that of *C. matruchotii*. We hypothesize that this should also be true for any other adjacent H_2_O_2_-producing streptococcal species. Transcriptional responses of *S. mitis* to *C. matruchotii* are part of a separate ongoing study and are not reflected here.

*C. matruchotii* has been shown to only oxidize lactate aerobically and cannot grow on lactate as a sole carbon source anaerobically (Iwami et al., 1972). Of the 22 genes that *C. matruchotii* differentially expresses with *S. mitis* aerobically, three belong to the *lutABC* operon which encodes lactate catabolism genes (Chai et al., 2009). We generated a deletion of the *lut* operon in *C. matruchotii* and found that it could only modestly oxidize lactate (Fig. S2) presumably due to reversible reaction(s) by any/all of 3 other L-lactate dehydrogenases that it encodes. We originally hypothesized that removal of lactate would benefit *S. mitis* by neutralizing the local pH. We tested this by addition of copious amounts of MOPS buffer but observed no significant changes in growth yield vs medium lacking MOPS. Using the pH indicator phenol red, we were unable to observe acidification around cocultured cells in the presence of additional buffer. Unexpectedly, these data suggest that pH modulation is not a factor in coculture growth yield benefit.

Alternatively, *C. matruchotii* may aid *S. mitis* by the removal of lactate itself and in doing so allow for more glucose fermentation by preventing feedback inhibition. This is further supported by total loss of coculture enhancement of *S. mitis* growth yield observed anaerobically (Fig. 1B). Lactate removal was linked to increased *S. mitis* growth in our limited glucose experiment which forced *C. matruchotii* to rely on lactate produced by *S. mitis* for full growth yield. We saw a significant decrease of *C. matruchotii ΔlutABC* when in coculture with *S. mitis* on limiting glucose media when compared to monoculture and a similar decrease in *S. mitis* yield (Fig. 2). This shows that *C. matruchotii* cannot compete for glucose when in competition with *S. mitis* and likely depends on cross-feeding of lactate when they are in direct proximity.

Using SECM we were able to directly quantify lactate production by *S. mitis* and its oxidation by adjacent *C. matruchotii* in real time (Fig. 5) indicating a sharp decrease in lactate concentration between individual cells. To the best of our knowledge this is the first observation of metabolite exchange between individual bacterial cells by SECM. We believe that existing ‘corncob’ configurations observed *in vivo* (Mark Welch et al., 2016; Morillo-lopez et al., 2021) should easily be able to consistently remove lactate from their immediate area. This would allow streptococcal metabolism to continue without inhibition while eliminating a source of acid stress to the host and other adjacent microbiota. This observation supports a mechanism whereby the interaction between these two commensals may contribute to the lack of cariogenic activity in a healthy oral biofilm.

This study has described two mechanisms of interaction between bacteria that exist in direct contact *in vivo*. Using a reductionist approach, we were able to ascertain how each mechanism contributed to fitness of both organisms. Advantages provided to each species when these mechanisms are intact also reflect the positional arrangement of these species *in vivo* as anaerobic conditions would not allow for favorable interactions to occur. Additionally, we were able to demonstrate real-time metabolite exchange between these species at sub-micron distances, indicating that crossfeeding between these organisms is likely occurring between them *in vivo*. These interactions reveal one way by which structural orientation and species composition between commensals may contribute to host health and potentially be one way by which a healthy biofilm composition is maintained *in vivo*.

## Acknowledgments

We thank Janet Atoyan and the RI-EPSCOR sequencing facility at URI for sequence generation, Jonathan Livny and the Microbial ‘Omics Core and Genomics Platform for their help with RNASeq library sequencing and guidance on experimental design, other members of the Ramsey lab and the Annual Mark Wilson conference attendees for many valuable suggestions and discussion.

## Author Contributions

MR and JK designed research; EA, SP, and SE performed research, EA, SP, JK and MR wrote the paper.

