## Supplemental data and methods for "*Corynebacterium matruchotii* fitness enhancement of adjacent streptococci by multiple mechanisms"

**Supplemental Materials**

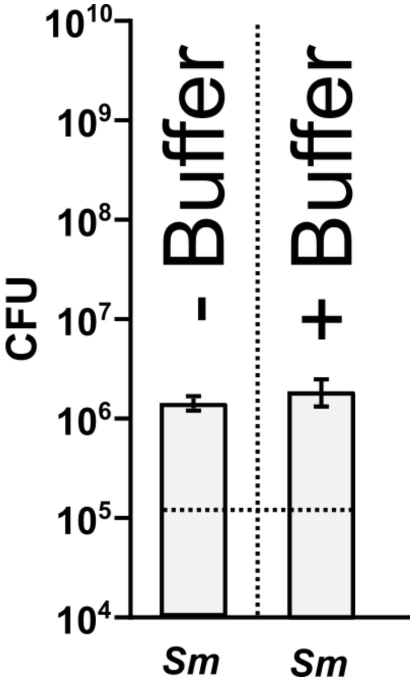

**FIG S1.** *S. mitis* (*Sm*) monoculture unaffected by the addition of 50mM MOPS to BHI-YE. *S. mitis* had a starting 1.4x10$^5$. Data are mean CFU counts with error bars indicating standard deviation for n≥3.

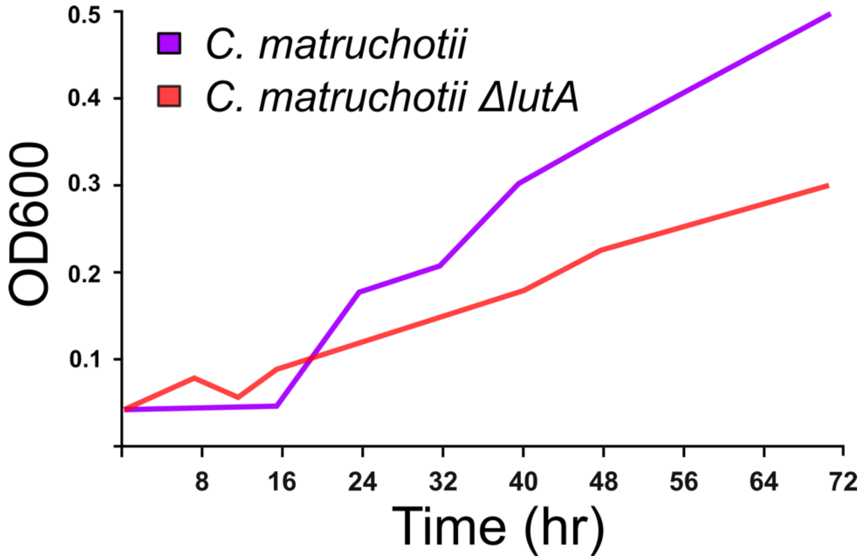

**FIG S2.** Growth curve of *C. matruchotii* and *C. matruchotii ΔlutA* grown on defined media containing lactate as the sole carbon source. Data are $OD_{600}$ readings for n = 1.

### Supplemental Methods and Results of SECM analysis

#### Fabrication of Submicrosized Pipet Electrodes

A quartz capillary (outer diameter 1 mm, inner diameter 0.7mm, length 10cm, Sutter instrument) was cleaned by blowing with compressed air, and pulled with $CO_2$ laser puller (Model P 2000-2035, Sutter Instrument) with parameters (heat = 720, filament = 4, velocity = 30, delay = 130 and pull = 110) giving c.a. 800 nm inner diameter. As-pulled pipets were placed inside UV plasma cleaner (PDC-32G, Harrick Plasma), and the chamber was degassed first, then filled with Ar gas as a source of plasma. As-pulled pipets were then cleaned by UV plasms for 3 minutes and again degassed for 10 minutes. Then, cleaned submicrosized pipets were dried for 1.5 hr under pressure below 200 mTorr inside a mini desiccator, which is placed inside a large glove bag filled with dry $N_2$ gas to maintain a constant relative humidity, 16 % at 20 $^0$C. Subsequently, silanization was done by chemical vapor deposition method with introducing 50 μL of N, N-dimethyltrimethylsilylamine to the desiccator, and left for 50 minutes under a constant relative humidity of 16 % at 20 $^0$C. Finally, the vacuum was applied to the desiccator to remove extra silanizing reagents for 5~10 min. Submicrozied pipets were then filled with 10 μl of 1,2-dichloroethane containing 0.1 M of tetrakis(pentafluorophenyl) borate and tetradodecyl-ammonium ions as supporting electrolytes. An electrochemically etched nickel/copper wire was inserted inside a submicrosized pipet as an inner reference electrode, and was immobilized for electrochemical measurements. The potential is applied between a submicropipet tip (a Ni/Cu inner reference electrode) and a Pt wire quasi reference electrode during SECM measurements.

#### Solution preparation for SECM measurements

Tetraethylammonium chloride ($TEA^+Cl^-$, Sigma Aldrich) solution of 1mM concentration was made in freshly prepared oral bacterial growth media of ~pH 7.5, where $TEA^+$ was used as a probe ion. The solution was then filtered with a syringe filter (0.1 μm filter unit, SLVV033RS, Duropore PVDF membrane, MILLEX VV). The SECM cell made with Teflon and glass was cleaned in piranha solution, and thoroughly rinsed with nanopure water (18.2 MΩ·cm, TOC 2 ppb; Milli-Q Integral 5 system, MIllipore).

#### NanoSECM measurements

Here, we used our home-built nanoSECM for measuring approach curves and imaging at single bacterial cells. Before approaching a submicrosized pipet tip to the bacteria, we measured cyclic voltammograms with tetraethylammonium ($TEA^+$, Sigma Aldrich) ion and estimate a steady state current in bulk solution ($i_{T,\infty}$) to confirm that nanopipet is working with a reasonable size. Then, our pipet was held c.a. 100 − 200 μm above the glass substrate with the help of a video microscope (Caltex VZM 400 lens with INFINITY2-1R 1.4 Megapixel USB 2.0 Microscopy Camera CCD) and a lockable micropositioner (DM-25L, Newport). We approached our pipet tip to the bacterial sample with z -piezo by measuring the steady-state current as a function of the distance between a pipet tip and glass substrate, and record an SECM approach curve. When the z-piezo moved 50 μm (maximum travel distance for piezo, P-620.ZCL, PI (Physik Instrumente), German) down at 50 nm/s, it was completely withdrawn and manually approached 50 μm with a lockable micropositioner. Then, the pipet tip was approached with z- piezo until it showed a sharp decrease in current, i.e., feedback current response. Once we observed a foot of feedback current response, the pipet tip was withdrawn 6 μm above, and approached at a slower rate i.e., 10 nm/s (1 nm step size with 100 ms incremental time) until our steady-state current decreased to c.a. 93 − 90 % of the $i_{T,\infty}$ , which corresponds to a distance of c.a. 2 times of tip radius from the substrate. Subsequently, a tip was further withdrawn 2 μm considering a height of bacteria. Here we approached only 93% of $i_{T,\infty}$ to prevent the crash with possible clumps of bacteria in coculture. Here, SECM imaging was done at the constant height mode. A pipet tip was raster scanned along *x*- and *y*-axis at 1 μm/s (i.e., 100 nm/100 ms) in the region of 25 μm × 25 μm to get a topographical image of the bacteria. After topographical imaging of the bacteria with respect to $TEA^+$, a pipet tip was brought to the original starting point. We again withdrew z-piezo 6 μm above, approached again at 10 nm/s (i.e., 1 nm/100 ms) until we reached 90 − 93 % of $i_{T,\infty}$, and withdrew 2 μm just right before imaging similar to previous topographic imaging step. A pipet tip potential is switched to a potential to sense lactate ion transfer (0.45 V more positive than $E_{1/2}$ of $TEA^+$ ion transfer, shown in Fig. S3B). Continuously, a tip was raster scanned in the same region of 25 μm × 25 μm to real time monitor lactate production/consumption by bacteria in situ.

**Direct, electrochemical detection of $TEA^+$ and Lactate ion transfer by a submicropipet-supported interface between two immiscible electrolyte solutions (ITIES)**

To investigate lactate production and consumption by bacteria as well as the topography of bacterial cells, a submicropipet-supported ITIES was employed. With this submicrotip, an etched Ni/Cu electrode in the internal organic electrolyte exerts a bias across the submicroscale liquid/liquid interface against an electrode in the aqueous solution (Fig. S3A) to yield the amperometric tip current based on the selective interfacial transfer of a small probe ion (Puri and Kim, 2019). A quartz submicropipet is filled with the electrolyte solution of 1,2-dichloroethane (DCE) to detect tetraethylammonium ($TEA^+$) as a probe ion, and lactate as a metabolite ion produced or consumed by bacteria. Herein, $TEA^+$ ion transfer (IT) is monitored at negative potential as a probe to study topography of bacterial sample, while lactate is sensed at positive potential to in situ monitor the chemical interaction between bacteria (Fig. S3B and S3C). The coculture of *C. matruchotii* and *S. mitis* is immobilized over a poly L-lysine coated slide glass plate, and studied by scanning or approaching a 800 nm-diameter pipet tip over the bacteria (Fig. S3C). Under negative potential applied to a submicrotip, the current response based on $TEA^+$ IT is lowered as the tip moves laterally toward the bacteria, which hinders the diffusion of the probe ion to the pipet tip. The tip current is recovered over the glass substrate (Fig. S3D), thereby probing topography of given bacterial sample. Once the tip potential is switched to more positive than $TEA^+$ IT (0.45 V more positive than $E_{1/2}$ of $TEA^+$ IT) and induces lactate IT, the tip directly senses lactate, thus leading to enhanced or lowered current response depending on production of consumption of lactate by bacteria, respectively in real time. Notably, a submicrometer-sized tip enables the real time study of chemical interactions between bacteria at a single cell level.

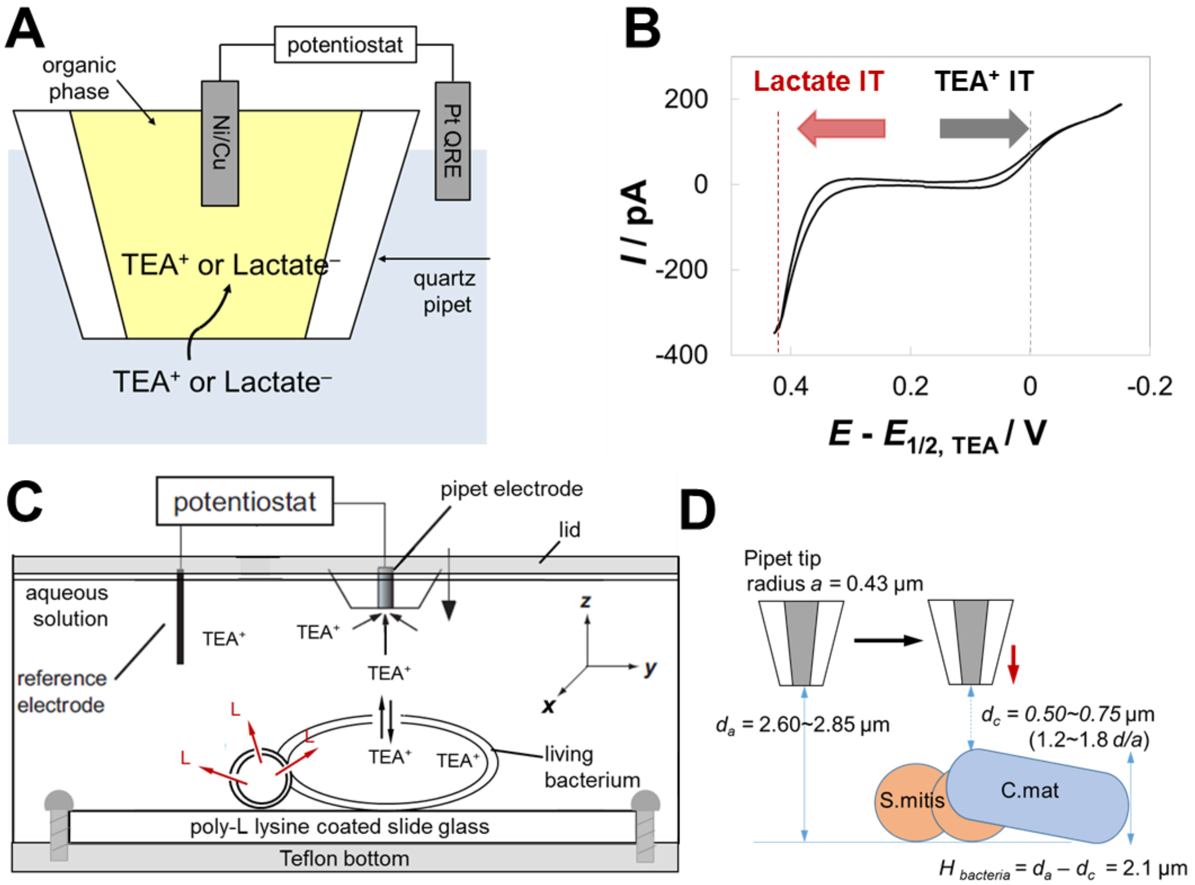

**Figure S3.** (A) Illustrated scheme of a submicropipet-supported ITIES to directly probe TEA$^+$ or lactate IT, (B) Cyclic voltammograms of TEA$^+$ and lactate ITs in buffer solution containing 1 mM TEA$^+$ and 2 mM lactate (background subtracted, scan rate = 25 mV/s), (C) Schematic of the SECM setup with living coculture bacterial samples immobilized on poly-L lysine coated slide glass, (D) Constant-height SECM imaging (black arrow) or SECM approach curve (red arrow) by a submicropipet tip scanned or vertically approached over *S. mitis* and *C. matruchotii* coculture, respectively.

**Quantitative analysis of the permeability of bacterial membranes, and local lactate concentration produced by *S. mitis* and consumed by *C. matruchotii***

After obtaining SECM images, an accurate position of bacteria could be specified. Subsequently, SECM approach curves were measured directly above *S. mitis* and *C. matruchotii*. The tip approached from the bulk solution while recording the steady-state tip current response for $TEA^+$ IT until contacting the surface of bacterial cell. This resulted in decrease in current as a function of normalized distance to the tip radius (*d/a*) with a sudden spike as a sign of contact between the tip and bacterial membrane (blue solid curves in Fig. S4A and B). The intrinsic permeability of both *S. mitis* and *C. matruchotii* membrane to $TEA^+$ ($k$ = 1.0 (± 0.1) × $10^{-3}$ cm/s) was determined by fitting experimental approach curves with a finite element simulation of a two phase SECM problem as described in detail elsewhere (blue open circles in Fig. S4A and B) (2.3). $TEA^+$ was used as a lieu of lactate for determining permeability for two reasons i.e., the same charge amount (± 1) and similar diffusion coefficients of ~6 × $10^{-6}$ $cm^2$/s. The approach curves over bacteria are nearly identical to the negative feedback approach curves obtained over an insulating glass substrate (grey solid curves in Fig. S4A and B), indicating that both bacterial membranes are almost impermeable to $TEA^+$.

With defining a permeability of *S. mitis* membrane as the determined $k$, an SECM approach curve over *S. mitis* was further simulated using a two phase SECM problem as well, where tip current responses for lactate IT become higher over *S. mitis* due to *in situ* generation of lactate. Herein, enhanced tip currents to 166 % of $i_{T,\infty}$ for lactate IT obtained at 1.20 d/a during SECM imaging (in Fig. 5C and 5E) are used as a criterion to verify the theoretically simulated approach curve. The resulting approach curve is depicted in Fig. S4C, where tip currents increase as the tip approaches over *S. mitis*, and reach 1.66 fold of $i_{T,\infty}$ at 1.2 *d/a*. The corresponding concentration profile of lactate near *S. mitis* and a pipet tip is shown in Fig. S4D, where the concentration of lactate produced by *S. mitis* is estimated as 0.52 mM very near the bacterial surface. This localized concentration of lactate is two times higher than that in bulk solutions resulting from the diffusion of lactate produced by other ensemble of *S. mitis* cells.

Notably, in approach curves over *C. matruchotii* (Fig. S4A), 80 % of $i_{T,\infty}$ at 1.20 *d/a* is consistent with tip current responses over *C. matruchotii* during constant-height SECM imaging based on lactate IT (~80 % current responses of $i_{T,\infty}$ over *C. matruchotii* in Fig. 5C and E). This result indicates that *C. matruchotii* efficiently depletes localized lactate produced by *S. mitis* in its proximity, thus tip current responses above *C. matruchotii* are not affected by localized lactate produced by *S. mitis* but controlled solely by its intrinsic membrane permeability and topography. Accordingly, tip current response based on lactate IT is lowered as the tip moves laterally toward *C. matruchotii*, which hinders the diffusion of the lactate ion pre-existing in bulk solution to the pipet tip.

In summary, this work demonstrates the high significance and power of SECM, which enabled us not only to *in situ* monitor the chemical communication between two commensal bacteria, *S. mitis* and *C. matruchotii* by imaging, but also to quantify the lactate production by *S. mitis* and consumption by *C. matruchotii* in real time at submicron resolution.

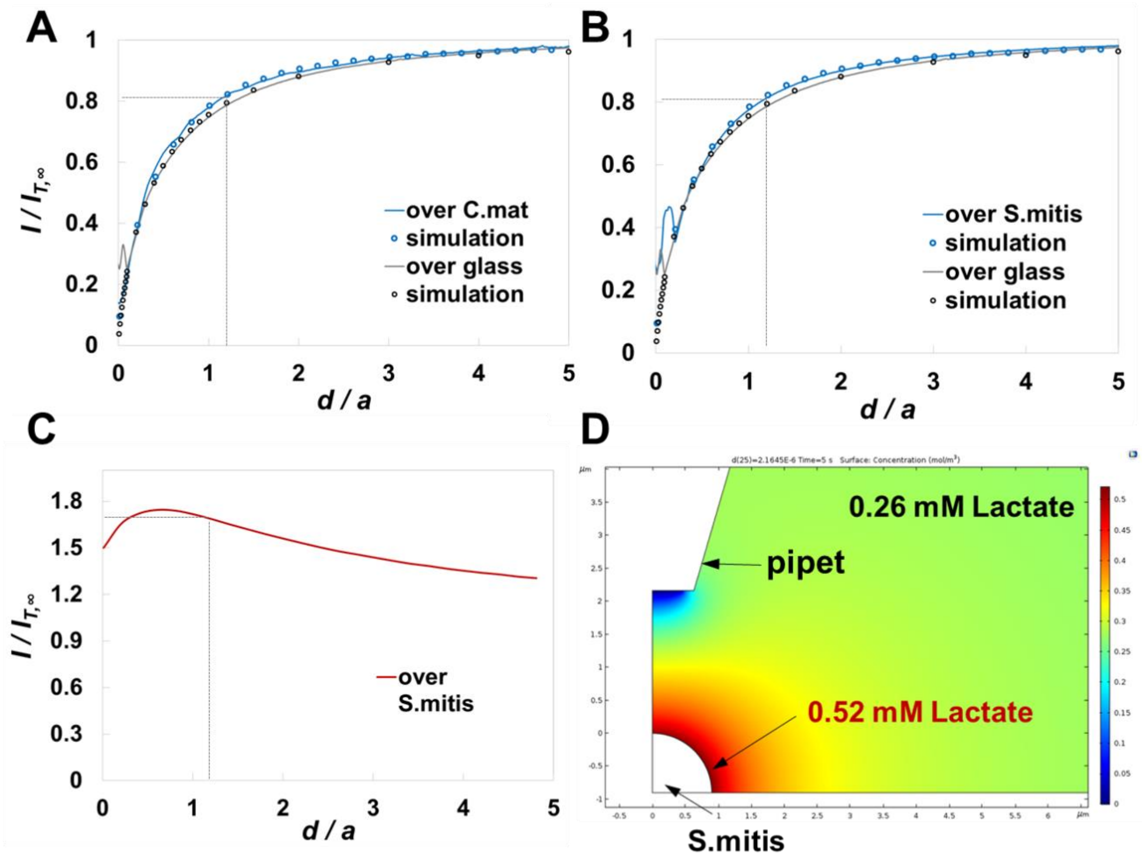

**Figure S4.** SECM approach curves based on $TEA^+$ IT (A) over *C. matruchotii* (blue solid line) and an insulating glass substrate (grey solid line), and (B) over *S. mitis* (blue solid line) and an insulating glass substrate (grey solid line). Experimental curves (blue solid lines) are compared with theoretically simulated curves (blue open circles) with permeability, $k$ = 1.0 (± 0.1) × $10^{-3}$ cm/s. Theoretically simulated negative feedback approach curves are shown as black open circles. (C) Simulated SECM approach curve based on lactate IT over S. mitis with considering determined $k$. (D) Concentration profile of lactate ions near *S. mitis* and a pipet tip.

**Table S1.** Lists of plasmids and strains used in study.

| Plasmids | Identifier | Host | Description |
|---|---|---|---|
| pEAKO2 | MR257 | DH5-alpha | Markerless clean deletion vector with sucrose counter-selection originated from pMRKO |
| pEA910 | MR317 | DH5-alpha | *lutABC* clean deletion vector utilizing pEAKO2 as parent vector |
| pEA400 | MR409 | DH5-alpha | *ftn* clean deletion vector utilizing pEAKO2 as parent vector |
| Species | Identifier | Strain | Description |
| *C. matruchotii* | MR127 | ATCC14266 | Wildtype strain |
| *C. matruchotii* | MR405 | *ΔlutABC* | *lutABC* operon clean deletion strain |
| *C. matruchotii* | MR414 | *Δftn* | *ftn* clean deletion strain |
| *S. mitis* | MR181 | ATCC49456 | Wildtype strain |
| *S. mitis* | MR291 | *ΔspxB* | *spxB* deletion strain |

**Table S2.** *C. matruchotii* differentially expressed genes when in coculture with *S. mitis* aerobically.

| fold_change | Gene ID | Gene Name |
|---|---|---|
| 6.77 | fig\|43768.439.peg.811 | FIG00544752: hypothetical protein |
| 4.37 | fig\|43768.439.peg.1549 | LutA Predicted L-lactate dehydrogenase Fe-S oxidoreductase subunit YkgE |
| 3.76 | fig\|43768.439.peg.1550 | LutB Predicted L-lactate dehydrogenase Iron-sulfur cluster-binding subunit YkgF |
| 3.20 | fig\|43768.439.peg.1551 | LutC Predicted L-lactate dehydrogenase hypothetical protein subunit YkgG |
| 2.96 | fig\|43768.439.peg.106 | hypothetical protein |
| 2.46 | fig\|43768.439.peg.2696 | LSU ribosomal protein L9p |
| 2.39 | fig\|43768.439.peg.491 | Bacterial non-heme ferritin (EC 1.16.3.2);Ontology_term |
| 2.10 | fig\|43768.439.peg.2766 | hypothetical protein |
| -2.02 | fig\|43768.439.peg.494 | Cytochrome c oxidase polypeptide I (EC 1.9.3.1);Ontology_term |
| -2.03 | fig\|43768.439.peg.1507 | ATP synthase epsilon chain (EC 3.6.3.14);Ontology_term |
| -2.09 | fig\|43768.439.peg.2518 | Proton/glutamate symporter @ Sodium/glutamate symporter |
| -2.11 | fig\|43768.439.peg.1692 | Enolase (EC 4.2.1.11);Ontology_term |
| -2.11 | fig\|43768.439.peg.1089 | hypothetical protein |
| -2.56 | fig\|43768.439.peg.1366 | PTS system beta-glucoside-specific IIB component / PTS system beta-glucoside-specific IIC component / PTS system beta-glucoside-specific IIA component |
| -2.65 | fig\|43768.439.peg.936 | Site-specific tyrosine recombinase XerC |

| | | |
|---|---|---|
| -2.80 | fig\|43768.439.peg.349 | Phosphoribosylformylglycinamidine synthase synthetase subunit (EC 6.3.5.3);Ontology_term |
| -3.00 | fig\|43768.439.peg.1196 | Iron-sulfur cluster regulator SufR |
| -4.10 | fig\|43768.439.peg.1202 | PaaD-like protein (DUF59) involved in Fe-S cluster assembly |
| -4.33 | fig\|43768.439.peg.1198 | Iron-sulfur cluster assembly protein SufD |
| -6.02 | fig\|43768.439.peg.1200 | Cysteine desulfurase (EC 2.8.1.7) > SufS;Ontology_term |
| -6.05 | fig\|43768.439.peg.1199 | Iron-sulfur cluster assembly ATPase protein SufC |

**Table S3.** *S. mitis* differentially expressed genes when in coculture with *C. matruchotii* aerobically.

| Fold Change | Gene ID | Gene Name |
|---|---|---|
| 75.83 | fig\|6666666.573673.peg.500 | hypothetical protein |
| 63.04 | fig\|6666666.573673.peg.1700 | Ribose ABC transporter%2C substrate-binding protein RbsB (TC 3.A.1.2.1) |
| 41.27 | fig\|6666666.573673.peg.601 | Zn-dependent alcohol dehydrogenases and related dehydrogenases |
| 40.84 | fig\|6666666.573673.peg.498 | hypothetical protein |
| 38.69 | fig\|6666666.573673.peg.503 | Putative protease |
| 34.00 | fig\|6666666.573673.peg.1704 | D-ribose pyranase (EC 5.4.99.62);Ontology_term |
| 29.34 | fig\|6666666.573673.peg.318 | N-acetylmannosamine-6-phosphate 2-epimerase (EC 5.1.3.9);Ontology_term |
| 28.98 | fig\|6666666.573673.peg.501 | Bacteriocin immunity protein BlpL |
| 22.72 | fig\|6666666.573673.peg.1703 | Ribose ABC transporter%2C ATP-binding protein RbsA (TC 3.A.1.2.1) |
| 22.22 | fig\|6666666.573673.peg.1705 | Ribokinase (EC 2.7.1.15);Ontology_term |
| 20.97 | fig\|6666666.573673.peg.497 | hypothetical protein |
| 20.66 | fig\|6666666.573673.peg.1699 | MutT/Nudix family protein |
| 17.99 | fig\|6666666.573673.peg.640 | hypothetical protein |
| 17.92 | fig\|6666666.573673.peg.1701 | Ribose ABC transporter%2C permease protein RbsC (TC 3.A.1.2.1) |
| 16.01 | fig\|6666666.573673.peg.260 | Transcriptional regulator SpxA1 |
| 14.51 | fig\|6666666.573673.peg.79 | UPF0223 protein YktA |
| 13.71 | fig\|6666666.573673.peg.80 | Transcriptional regulator SpxA1 |
| 13.27 | fig\|6666666.573673.peg.499 | FIG01119121: hypothetical protein |
| 13.08 | fig\|6666666.573673.peg.813 | Surface protein PspC |
| 12.52 | fig\|6666666.573673.peg.547 | Alcohol dehydrogenase (EC 1.1.1.1);Ontology_term |
| 12.32 | fig\|6666666.573673.peg.502 | BlpZ protein%2C fusion |
| 11.19 | fig\|6666666.573673.peg.814 | Surface protein PspC |

| | | |
|---|---|---|
| 10.61 | fig\|6666666.573673.peg.316 | ABC transporter%2C predicted N-acetylneuraminate-binding protein |
| 9.79 | fig\|6666666.573673.peg.327 | hypothetical protein |
| 9.15 | fig\|6666666.573673.peg.326 | hypothetical protein |
| 8.69 | fig\|6666666.573673.peg.285 | Manganese ABC transporter%2C ATP-binding protein SitB |
| 8.31 | fig\|6666666.573673.peg.548 | PTS system%2C mannose-specific IIA component (EC 2.7.1.191) / PTS system%2C mannose-specific IIB component (EC 2.7.1.191);Ontology_term |
| 8.15 | fig\|6666666.573673.peg.895 | hypothetical protein |
| 7.57 | fig\|6666666.573673.peg.812 | Surface protein PspC |
| 7.54 | fig\|6666666.573673.peg.550 | PTS system%2C mannose-specific IID component |
| 7.40 | fig\|6666666.573673.peg.721 | hypothetical protein |
| 7.31 | fig\|6666666.573673.peg.1336 | Rhodanese-like domain protein |
| 7.02 | fig\|6666666.573673.peg.717 | Trans-acting positive regulator |
| 6.42 | fig\|6666666.573673.peg.842 | PTS system%2C galactosamine-specific IIA component |
| 6.14 | fig\|6666666.573673.peg.720 | putative membrane protein |
| 5.94 | fig\|6666666.573673.peg.1204 | Maltodextrin ABC transporter%2C ATP-binding protein MsmX |
| 5.79 | fig\|6666666.573673.peg.1312 | Two-component response regulator yesN%2C associated with MetSO reductase |
| 5.65 | fig\|6666666.573673.peg.1140 | Biotin carboxyl carrier protein of acetyl-CoA carboxylase |
| 5.61 | fig\|6666666.573673.peg.1122 | Ribonuclease HIII (EC 3.1.26.4);Ontology_term |
| 5.42 | fig\|6666666.573673.peg.1610 | Phosphoenolpyruvate-dihydroxyacetone phosphotransferase (EC 2.7.1.121)%2C subunit DhaM%3B DHA-specific IIA component;Ontology_term |
| 5.31 | fig\|6666666.573673.peg.645 | Maltodextrin ABC transporter%2C permease protein MdxG |
| 5.30 | fig\|6666666.573673.peg.680 | Putative adhesin SPy2174 |
| 5.28 | fig\|6666666.573673.peg.644 | Maltodextrin ABC transporter%2C permease protein MdxF |
| 5.17 | fig\|6666666.573673.peg.841 | PTS system%2C galactosamine-specific IID component |
| 5.07 | fig\|6666666.573673.peg.372 | beta-fructofuranosidase (EC 3.2.1.26);Ontology_term |
| 5.04 | fig\|6666666.573673.peg.317 | PTS system%2C N-acetylmannosamine-specific IIC component / PTS system%2C N-acetylmannosamine-specific IIB component |
| 5.02 | fig\|6666666.573673.peg.549 | PTS system%2C mannose-specific IIC component |
| 5.00 | fig\|6666666.573673.peg.643 | Maltodextrin ABC transporter%2C substrate-binding protein MdxE |

| | | |
|---|---|---|
| 5.00 | fig\|6666666.573673.peg.802 | Merozoite surface protein 3 alpha |
| 4.98 | fig\|6666666.573673.peg.681 | Putative uncharacterized protein spr0086 |
| 4.89 | fig\|6666666.573673.peg.178 | PTS system%2C maltose and glucose-specific IIC component / PTS system%2C maltose and glucose-specific IIB component (EC 2.7.1.208) / PTS system%2C maltose and glucose-specific IIA component;Ontology_term |
| 4.80 | fig\|6666666.573673.peg.679 | Uncharacterized membrane protein SPy2173 |
| 4.66 | fig\|6666666.573673.peg.742 | Cysteine synthase (EC 2.5.1.47);Ontology_term |
| 4.50 | fig\|6666666.573673.peg.1380 | Uncharacterized membrane spanning protein%2C contains VanZ-like domain |
| 4.49 | fig\|6666666.573673.peg.973 | Ferric iron ABC transporter%2C ATP-binding protein |
| 4.43 | fig\|6666666.573673.peg.991 | Pullulanase (EC 3.2.1.41);Ontology_term |
| 4.31 | fig\|6666666.573673.peg.972 | Ferric iron ABC transporter%2C permease protein |
| 4.25 | fig\|6666666.573673.peg.284 | Manganese ABC transporter%2C ATP-binding protein SitB |
| 4.19 | fig\|6666666.573673.peg.198 | beta-galactosidase (EC 3.2.1.23);Ontology_term |
| 4.09 | fig\|6666666.573673.peg.1153 | hypothetical protein |
| 4.01 | fig\|6666666.573673.peg.1012 | Pyruvate formate-lyase (EC 2.3.1.54);Ontology_term |
| 3.97 | fig\|6666666.573673.peg.678 | Transcriptional regulator SPy2172%2C PadR family |
| 3.94 | fig\|6666666.573673.peg.1262 | hypothetical protein |
| 3.91 | fig\|6666666.573673.peg.581 | Acetaldehyde dehydrogenase (EC 1.2.1.10) / Alcohol dehydrogenase (EC 1.1.1.1);Ontology_term |
| 3.84 | fig\|6666666.573673.peg.715 | Alpha-glycerophosphate oxidase (EC 1.1.3.21);Ontology_term |
| 3.84 | fig\|6666666.573673.peg.287 | Manganese ABC transporter%2C periplasmic-binding protein SitA |
| 3.74 | fig\|6666666.573673.peg.974 | Ferric iron ABC transporter%2C iron-binding protein |
| 3.73 | fig\|6666666.573673.peg.288 | Thiol peroxidase%2C Tpx-type (EC 1.11.1.15);Ontology_term |
| 3.73 | fig\|6666666.573673.peg.559 | BOX elements |
| 3.65 | fig\|6666666.573673.peg.1706 | Ribose operon repressor |
| 3.65 | fig\|6666666.573673.peg.1612 | Phosphoenolpyruvate-dihydroxyacetone phosphotransferase (EC 2.7.1.121)%2C dihydroxyacetone binding subunit DhaK;Ontology_term |
| 3.59 | fig\|6666666.573673.peg.1328 | Methionine biosynthesis and transport regulator MtaR%2C LysR family |
| 3.58 | fig\|6666666.573673.peg.52 | ABC transporter%2C permease protein |
| 3.56 | fig\|6666666.573673.peg.310 | N-acetylmannosamine kinase (EC 2.7.1.60);Ontology_term |

| | | |
|---|---|---|
| 3.56 | fig\|6666666.573673.peg.177 | Exodeoxyribonuclease III (EC 3.1.11.2);Ontology_term |
| 3.53 | fig\|6666666.573673.peg.716 | Glycerol kinase (EC 2.7.1.30);Ontology_term |
| 3.47 | fig\|6666666.573673.peg.838 | beta-galactosidase (EC 3.2.1.23);Ontology_term |
| 3.47 | fig\|6666666.573673.peg.637 | Lead%2C cadmium%2C zinc and mercury transporting ATPase (EC 3.6.3.3) (EC 3.6.3.5)%3B Copper-translocating P-type ATPase (EC 3.6.3.4);Ontology_term |
| 3.36 | fig\|6666666.573673.peg.1614 | dihydroxyacetone kinase family protein |
| 3.34 | fig\|6666666.573673.peg.714 | Glycerol uptake facilitator protein |
| 3.12 | fig\|6666666.573673.peg.289 | ABC transporter%2C FliJ domain and fused permease subunit |
| 3.03 | fig\|6666666.573673.peg.1364 | Hydroxymethylpyrimidine kinase (EC 2.7.1.49) @ Hydroxymethylpyrimidine phosphate kinase ThiD (EC 2.7.4.7);Ontology_term |
| 2.98 | fig\|6666666.573673.peg.1441 | hypothetical protein |
| 2.86 | fig\|6666666.573673.peg.228 | Glycine/D-amino acid oxidases family |
| 2.77 | fig\|6666666.573673.peg.1509 | hypothetical protein |
| 2.66 | fig\|6666666.573673.peg.634 | Lead%2C cadmium%2C zinc and mercury transporting ATPase (EC 3.6.3.3) (EC 3.6.3.5)%3B Copper-translocating P-type ATPase (EC 3.6.3.4);Ontology_term |
| 2.60 | fig\|6666666.573673.peg.381 | General stress protein%2C Gls24 family |
| 2.56 | fig\|6666666.573673.peg.1081 | Exopolysaccharide biosynthesis transcriptional activator EpsA |
| 2.55 | fig\|6666666.573673.peg.875 | DUF402 family nucleoside diphosphatase |
| 2.53 | fig\|6666666.573673.peg.737 | Ribosome hibernation promoting factor Hpf |
| 2.49 | fig\|6666666.573673.peg.801 | Surface protein PspC |
| 2.46 | fig\|6666666.573673.peg.1104 | Cell division protein GpsB%2C coordinates the switch between cylindrical and septal cell wall synthesis by re-localization of PBP1 |
| 2.46 | fig\|6666666.573673.peg.414 | Glycine betaine ABC transport system%2C permease protein OpuAB / Glycine betaine ABC transport system%2C glycine betaine-binding protein OpuAC |
| 2.24 | fig\|6666666.573673.peg.1210 | Single-stranded-DNA-specific exonuclease RecJ |
| 2.23 | fig\|6666666.573673.peg.475 | Endo-beta-N-acetylglucosaminidase (EC 3.2.1.96);Ontology_term |
| 2.00 | fig\|6666666.573673.peg.538 | Catabolite control protein A |
| -2.01 | fig\|6666666.573673.peg.1411 | Two component system sensor histidine kinase CiaH (EC 2.7.3.-);Ontology_term |
| -2.03 | fig\|6666666.573673.peg.1714 | Glucose-6-phosphate 1-dehydrogenase (EC 1.1.1.49);Ontology_term |
| -2.05 | fig\|6666666.573673.peg.1144 | Acetyl-coenzyme A carboxyl transferase alpha chain (EC 6.4.1.2);Ontology_term |

| | | |
|---|---|---|
| -2.08 | fig\|6666666.573673.peg.1027 | ABC transporter membrane-spanning permease - Na+ export |
| -2.11 | fig\|6666666.573673.peg.432 | FIG005935: membrane protein |
| -2.15 | fig\|6666666.573673.peg.785 | Cell division-associated%2C ATP-dependent zinc metalloprotease FtsH |
| -2.24 | fig\|6666666.573673.peg.487 | Intracellular protease |
| -2.27 | fig\|6666666.573673.peg.1686 | Cysteine desulfurase (EC 2.8.1.7);Ontology_term |
| -2.29 | fig\|6666666.573673.peg.726 | ATP-dependent Clp protease%2C ATP-binding subunit ClpC |
| -2.33 | fig\|6666666.573673.peg.1622 | Ribonucleotide reductase of class Ib (aerobic)%2C beta subunit (EC 1.17.4.1);Ontology_term |
| -2.42 | fig\|6666666.573673.peg.1264 | M protein trans-acting positive regulator (Mga) |
| -2.43 | fig\|6666666.573673.peg.768 | Serine protease%2C DegP/HtrA%2C do-like (EC 3.4.21.-);Ontology_term |
| -2.43 | fig\|6666666.573673.peg.151 | Hemolysin III |
| -2.44 | fig\|6666666.573673.peg.989 | Glutamine--fructose-6-phosphate aminotransferase [isomerizing] (EC 2.6.1.16);Ontology_term |
| -2.57 | fig\|6666666.573673.peg.350 | Aquaporin Z |
| -2.58 | fig\|6666666.573673.peg.1541 | Xaa-Pro dipeptidyl-peptidase (EC 3.4.14.11);Ontology_term |
| -2.59 | fig\|6666666.573673.peg.1764 | LSU ribosomal protein L19p |
| -2.64 | fig\|6666666.573673.peg.485 | Chaperone protein DnaK |
| -2.66 | fig\|6666666.573673.peg.483 | Heat-inducible transcription repressor HrcA |
| -2.68 | fig\|6666666.573673.peg.1621 | Ribonucleotide reductase of class Ib (aerobic)%2C alpha subunit (EC 1.17.4.1);Ontology_term |
| -2.69 | fig\|6666666.573673.peg.1569 | UPF0703 protein YcgQ |
| -2.69 | fig\|6666666.573673.peg.484 | Heat shock protein GrpE |
| -2.88 | fig\|6666666.573673.peg.1559 | hypothetical protein |
| -2.88 | fig\|6666666.573673.peg.710 | D-alanyl transfer protein DltB |
| -2.91 | fig\|6666666.573673.peg.667 | tRNA-specific 2-thiouridylase MnmA (EC 2.8.1.13);Ontology_term |
| -2.97 | fig\|6666666.573673.peg.1524 | Putative parvulin type peptidyl-prolyl isomerase%2C similarity with PrsA foldase |
| -3.00 | fig\|6666666.573673.peg.1273 | Cell wall surface anchor family protein |
| -3.05 | fig\|6666666.573673.peg.1557 | 23S rRNA (uracil(1939)-C(5))-methyltransferase (EC 2.1.1.190);Ontology_term |
| -3.22 | fig\|6666666.573673.peg.215 | Putative amidotransferase similar to cobyric acid synthase |
| -3.23 | fig\|6666666.573673.peg.1681 | Similar to ribosomal large subunit pseudouridine synthase D%2C Bacillus subtilis YjbO type |
| -3.30 | fig\|6666666.573673.peg.765 | Histidine kinase of the competence regulon ComD |

| | | |
|---|---|---|
| -3.32 | fig\|6666666.573673.peg.1393 | hypothetical protein |
| -3.35 | fig\|6666666.573673.peg.213 | Putative Dihydrolipoamide dehydrogenase (EC 1.8.1.4)%3B Mercuric ion reductase (EC 1.16.1.1)%3B PF00070 family%2C FAD-dependent NAD(P)-disulphide oxidoreductase;Ontology_term |
| -3.58 | fig\|6666666.573673.peg.664 | tRNA-5-carboxymethylaminomethyl-2-thiouridine(34) synthesis protein MnmG |
| -3.58 | fig\|6666666.573673.peg.846 | IgA1 protease (EC 3.4.24.13);Ontology_term |
| -3.64 | fig\|6666666.573673.peg.214 | proposed amino acid ligase found clustered with an amidotransferase |
| -3.68 | fig\|6666666.573673.peg.764 | Response regulator of the competence regulon ComE |
| -3.68 | fig\|6666666.573673.peg.749 | Transmembrane component of general energizing module of ECF transporters |
| -3.75 | fig\|6666666.573673.peg.891 | Cytochrome c-type biogenesis protein CcdA (DsbD analog) |
| -3.78 | fig\|6666666.573673.peg.1576 | Phage shock protein C%2C putative |
| -3.79 | fig\|6666666.573673.peg.925 | Mg(2+) transport ATPase%2C P-type (EC 3.6.3.2);Ontology_term |
| -4.17 | fig\|6666666.573673.peg.1692 | ATP-dependent DNA helicase UvrD/PcrA (EC 3.6.4.12);Ontology_term |
| -4.61 | fig\|6666666.573673.peg.1115 | Membrane protein LiaF(VraT)%2C specific inhibitor of LiaRS(VraRS) signaling pathway |
| -4.66 | fig\|6666666.573673.peg.823 | Competence-stimulating peptide ABC transporter ATP-binding protein ComA |
| -5.65 | fig\|6666666.573673.peg.1556 | Extracellular protein |
| -5.78 | fig\|6666666.573673.peg.1118 | DNA alkylation repair enzyme |
| -5.84 | fig\|6666666.573673.peg.1117 | Cell envelope stress response system LiaFSR%2C response regulator LiaR(VraR) |
| -6.09 | fig\|6666666.573673.peg.1116 | Cell envelope stress response system LiaFSR%2C sensor histidine kinase LiaS(VraS) |
| -7.66 | fig\|6666666.573673.peg.824 | Competence-stimulating peptide ABC transporter permease protein ComB |
